# Neural signatures of recollection are sensitive to memory quality and specific event features

**DOI:** 10.1101/2025.03.18.643924

**Authors:** Natalia Ladyka-Wojcik, Helen Schmidt, Rose A. Cooper, Maureen Ritchey

**Affiliations:** Department of Psychology & Neuroscience, Boston College, Chestnut Hill, MA, USA; Department of Psychology and Neuroscience, Temple University, Philadelphia, PA, USA

## Abstract

Episodic memories reflect a bound representation of multimodal features that can be recollected with varying levels of precision. Recent fMRI investigations have demonstrated that the precision and content of information retrieved from memory engage a network of posterior medial temporal and parietal regions co-activated with the hippocampus. Yet, comparatively little is known about how memory content and precision affect common neural signatures of memory captured by electroencephalography (EEG), where recollection has been associated with changes in event-related potential (ERP) and oscillatory measures of neural activity. Here, we used a multi-feature paradigm previously reported in Cooper & Ritchey (2019) with continuous measures of memory, in conjunction with scalp EEG, to characterize the content and quality of information that drives ERP and oscillatory markers of episodic memory. A common signature of memory retrieval in left posterior regions, called the late positive component (LPC), was sensitive to overall memory quality and also to precision of recollection for spatial features. Analysis of oscillatory markers during recollection revealed that alpha/beta desynchronization was modulated by overall memory quality and also by individual features in memory. Importantly, we found evidence of a relationship between these two neural markers of memory retrieval, suggesting that they may represent complementary aspects of the recollection experience. These findings demonstrate how time-sensitive and dynamic processes identified with EEG correspond to overall episodic recollection, and also to the retrieval of precise features in memory.

## Introduction

Recollecting a past event, such as a trip to the beach or a birthday party, involves reactivating multiple event features, including people, items, places, colors, and sounds, and representing those features as a coherent episode. Episodic recollection, compared to other forms of memory, engages a network of posterior medial temporal and parietal regions co-activated with the hippocampus, including the parahippocampal cortex, retrosplenial cortex, precuneus, medial prefrontal cortex, and angular gyrus (Addis et al., 2007; Benoit & Schacter, 2015; Ritchey & Cooper, 2020; Rugg & Vilberg, 2013; Spreng et al., 2009). Different aspects of recollection, such as memory precision and fidelity, have been shown to variably relate to activity in these regions (Nilakantan et al., 2018; Richter et al., 2016; Vilberg & Rugg, 2007) and greater integration of content in episodic representations corresponds to their increased functional interconnectivity (Cooper & Ritchey, 2019). Moreover, recollection has been tied to a set of unique neural signatures using electroencephalography (EEG; Addante et al., 2012; Hanslmayr et al., 2012; Rugg & Curran, 2007), but it remains unclear if these dynamic signatures reflect both the precision and multidimensional content of episodic memory. In this study, we investigated whether two previously identified EEG markers of recollection, via event-related potentials (ERPs) and oscillatory dynamics, are sensitive to the representational quality of recollective experience.

Recollective processes have most often been studied in both behavioral and fMRI research using binary measures of memory, such as successful versus unsuccessful retrieval of contextual information, or with subjective judgments of vividness. These methods cannot capture the objective precision or the representational content of memory, highlighting the importance of incorporating additional measures to assess the quality of memory. An approach that accounts for both the amount of information in memory (Vilberg & Rugg, 2007) and memory fidelity (Brady et al., 2013) is necessary to characterize the representational content of recollective experience. Consistent with this approach, some research has leveraged continuous measures of memory performance (see e.g., Harlow & Donaldson, 2013) to demonstrate behavioral and neural dissociations between retrieval success and memory precision. For example, studies quantifying memory resolution for item-location associations with a continuous measure have shown that retrieval precision varies on a trial-by-trial basis but remains relatively stable over time (Berens et al., 2020), whereas unsuccessful retrieval (i.e., forgetting) involves the loss of all associative information in memory (Richter et al., 2016; Vilberg & Rugg, 2007).

Precision in episodic retrieval for item-location associations seems to recruit parietal regions such as the angular gyrus, in contrast to memory success in the hippocampus proper (Richter et al., 2016). Moreover, whereas binary memory success demonstrates a degree of dependency across features (Horner & Burgess, 2013), the specific resolution of each feature in memory appears to be somewhat independent, further justifying the use of continuous measures that can capture multiple features in memory representations. Indeed, Cooper and Ritchey (2019) also found that integration of cortico-hippocampal networks across both anterior-temporal and posterior-medial regions of the brain is modulated by the multidimensional quality of episodic recollection, and that specific patterns of network integration are observable for memories with precise representations of scene perspective and item color. Leveraging the temporal resolution of EEG in conjunction with this type of continuous, multidimensional memory paradigm could help to pull apart the components of recollection and afford greater specificity in characterizing the temporal dynamics of memory retrieval.

Prior EEG studies have investigated the neural correlates of memory retrieval in two key ways: with ERPs and with time-frequency (i.e., oscillatory) analyses. Early investigations of ERPs revealed a distinct characteristic of memory retrieval termed the “late positive component” (LPC; Addante et al., 2012), or the left parietal old/new effect (for a review, see Vilberg & Rugg, 2007), wherein correct item retrieval is associated with an ERP modulation around 500-800 ms over posterior, particularly left, electrodes. Beyond item retrieval, the LPC is particularly sensitive to memory for pictures over words (Curran & Doyle, 2011) and correct source memory (Addante et al., 2012; Diana et al., 2011), indicating some correspondence to the retrieval of visual-perceptual information. The LPC also appears to be driven specifically by recollection as opposed to familiarity-based subjective judgments of memory (Curran, 2004; Duarte et al., 2004; Vilberg & Rugg, 2009). Additionally, studies suggest that the LPC reflects the amount of contextual information recalled and that it captures the reactivation of event information from memory. Specifically, Vilberg et al. (2006) reported that the LPC demonstrated greater amplitude when elicited by fully recollected visual items compared to partially recollected ones.

However, exactly how the LPC signature of recollection reflects the specific contents and precision of a reactivated memory is unclear. To our knowledge, only one study has implemented a continuous measure of recollection in conjunction with ERP analysis, quantifying the precision of memory for words in specific spatial locations (Murray et al., 2015). The authors found graded modulation of the LPC from high to low precision retrieval, but a total absence of this ERP signature for recollection failure. Whether graded sensitivity of the LPC to retrieval precision extends beyond spatial associations to other types of features in memory (e.g., color, sound, emotion, etc.) in a multidimensional context remains unclear. In short, the LPC could be characterized by sensitivity to memory fidelity for any given feature or could more broadly reflect the multidimensional quality of bound episodic representations.

A second signature of episodic memory revealed by EEG is oscillatory desynchronization, with desynchronization in the alpha and beta frequency bands being associated with recollection in contrast to forgetting (Hanslmayr et al., 2012). Specifically, the effect is believed to reflect the reactivation of event features and the amount or richness of information that can be retrieved. In support, a recent study found that alpha power 500 ms post-retrieval onset distinguished between item recognition and associative retrieval over left posterior electrodes, with the lowest alpha power for associated retrieval (Martín-Buro et al., 2020). Moreover, cortical desynchronization is thought to reactivate the specific contents of a memory, as evidenced by variation in the topography of alpha desynchronization with the kind of material recalled (Hanslmayr et al., 2016; Khader & Rösler, 2011). It has also been found to track the number of items recalled in memory (Fukuda & Woodman, 2017). Beyond binary or discrete measures of memory, the use of continuous measures of memory performance also reveals that alpha power is graded by the precision with which single features are remembered. For example, Sutterer et al. (2019) observed alpha-band desynchronization with precise memory retrieval for spatial locations. Yet, recollective experience in everyday life is not limited to single features and less is known about how exactly this oscillatory effect is modulated by high-fidelity retrieval of multiple features in memory. Alpha/beta desynchronization may be tied to the recollection of any given distinct features within an episodic memory, regardless of the relational structure of its constituent features. In this case, we would expect similar patterns of desynchronization tracking precision of memory for each specific feature, even when other features are not remembered precisely. For example, Griffiths et al. (2019) demonstrated that decreases in alpha/beta power during a retrieval task of visually-complex information tracked the fidelity of the stimulus-specific information in fMRI BOLD signal. Alternatively, this effect could be graded by the recollection quality of the overall episodic memory, reflecting multiple features bound into a coherent representation. Here, we would expect the magnitude of alpha/beta desynchronization to scale with overall memory quality, but this has yet to be tested using a continuous, multidimensional memory paradigm. Thus, an open question is whether alpha/beta desynchronization is sensitive to both the precision and multidimensional, relational content of episodic recollection.

In the current study, we asked to what extent two EEG signatures of recollection — the LPC and alpha/beta desynchronization — are modulated by the multidimensional quality of episodic memory and whether they are sensitive to the content of memory representations. To do so, we employed a task (Cooper & Ritchey, 2019) in which trial-unique items are presented in a color, selected from a continuous color spectrum, a specific scene within a 360-degree panorama, with an associated sound. This task allowed us to characterize the overall quality of episodic memory, in terms of the total number and specificity of features recalled, as well as memory for individual features. To preview the results, we found that the LPC was sensitive to overall memory quality, and reflected precise recollection of spatial location. Furthermore, we demonstrated that alpha/beta desynchronization during recollection was modulated by overall memory quality and also by specific features in memory. Finally, there was a relationship between memory-modulated activity in the LPC window and alpha/beta desynchronization during the later stages of retrieval. Overall, our results show how two EEG signatures of recollection dynamically capture the quality and content of precise, multidimensional memory recollection.

## Methods

### Participants

Data from 23 young adults (19 women, ages 18 - 22) were included in the final analyses, similar to the sample sizes of relevant EEG investigations of episodic memory (see e.g., Griffiths et al., 2019; Staudigl et al., 2010) and of a recent fMRI investigation using the same episodic memory task (Cooper & Ritchey, 2019). An additional ten participants completed the study but were excluded from analyses for the following reasons: (1) chance-level performance on the experimental task (*n* = 3); (2) high number of noisy channels throughout the experiment (*n* = 1); and (3) insufficient number of retrieval trials with good EEG data quality (*n* = 6; see EEG Data Processing). All participants had normal or corrected-to-normal vision and had no current diagnoses or history of psychological or neurological disorders. Informed consent was obtained from all participants. All procedures were approved by the Boston College Institutional Review Board, and participants received course credit or cash payment for their time.

### Experimental Design

The episodic memory task was designed and presented using PsychToolbox in MATLAB (Kleiner et al., 2007), and was identical to that used by our lab in a prior MRI study (Cooper & Ritchey, 2019). The stimuli included 144 objects selected from https://bradylab.ucsd.edu/stimuli.html as used in Brady et al. (2013), 12 emotional and neutral sounds selected from the International Affective Digitized Sounds (IADS) database (Bradley & Lang, 2007), and six panorama scenes selected from the SUN 360 database (http://3dvision.princeton.edu/projects/2012/SUN360/; Xiao et al., 2012).

To summarize, participants completed six independent encoding-retrieval blocks of 24 trials, for a total of 144 encoding trials and 144 retrieval trials. During encoding, participants viewed and were instructed to remember an item in a specific color, sampled from a 360-degree color spectrum, placed within a scene sampled from a 360-degree panoramic image. They also heard one of six negative sounds (e.g., a woman screaming) or one of six neutral sounds (e.g., a lawn mower) with each item, which they were asked to use to remember the object as a “bomb” or as “safe”, respectively. This instruction encouraged participants to integrate the objects and its associated features into a meaningful event (Figure 1A). Within a study block, each scene panorama was shown four times and each sound was encoded twice. All objects were trial unique. The object color and scene location values were pseudo-randomly selected with the constraint that objects associated with the same panorama within the same block should be at least 45 degrees apart in their color and location within the scene to minimize interference. During a subsequent memory test, participants were cued with one of the studied items presented in grayscale. First, they had four seconds to covertly remember as much as possible about the item’s color, scene location, and associated sound (Figure 1B). The EEG analyses focused on this four-second covert recall period, although EEG data was collected continuously throughout the experiment. Next, participants had two seconds to respond if the sound associated with that item was negative or neutral on a scale of 1 – 4 (1 = high confidence, negative; 2 = low confidence, negative; 3 = low confidence, neutral; 4 = high confidence, neutral). Participants then had six seconds to cycle through a circular color wheel and select the color they remembered the item to be during the earlier viewing period. Once they made their color selection, they had another six seconds to pan through the scenes within the 360-degree panorama to select the scene they remembered (Figure 1C). The order of the color and scene questions was counterbalanced across trials, and the order of trials was randomized for every participant and every encoding and retrieval phase. Please see Cooper & Ritchey (2019) for additional details of the task design.

**Figure 1.**
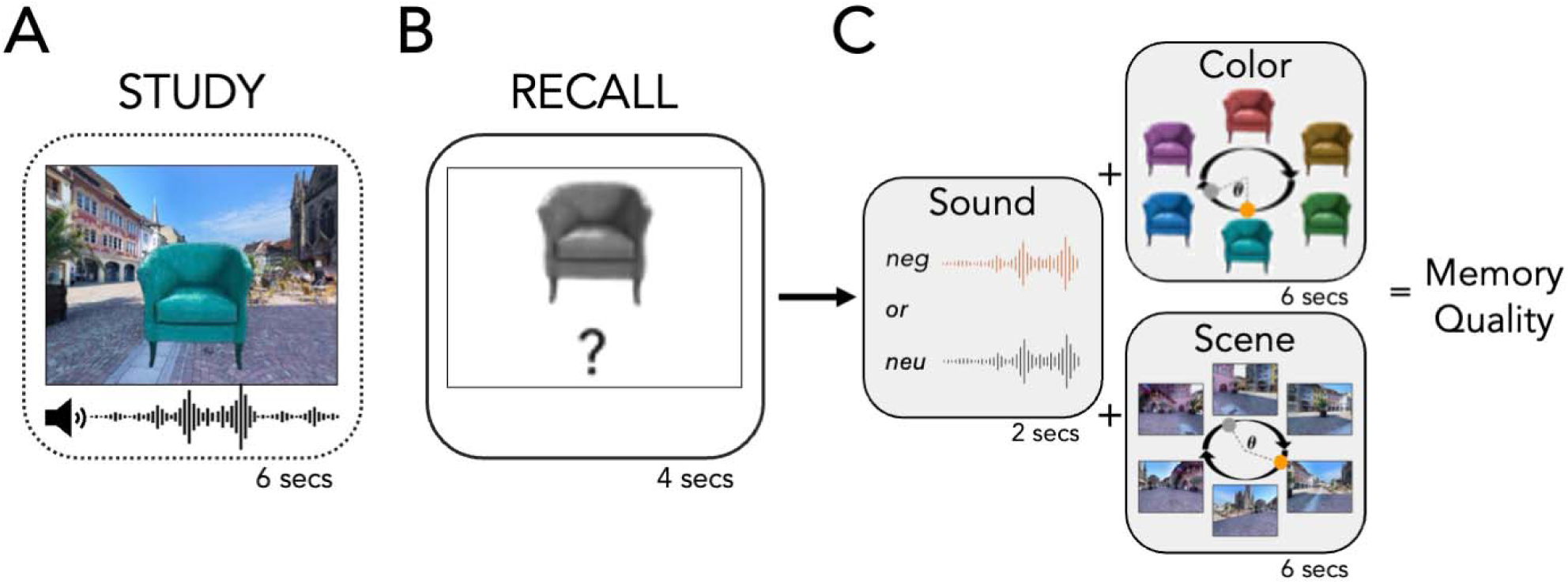
Behavioral measures of memory. (A) Participants studied items (dashed box) associated with three features – a color, a scene location, and an emotional sound. (B) In a subsequent memory recall test (solid box), participants were cued with a grayscale item and were asked to covertly recall all of the associated features for 4 seconds (gray boxes). This recall period is the focus of all EEG analyses. (C) In subsequent questions, we assessed memory for each feature separately, including a negative (neg) or neutral (neu) decision and confidence rating for the sound, and selection of a color and panorama scene perspective from 360 degrees spaces. Color and scene responses were measured in terms of error in degrees between the participant’s response (gray circle) and the correct value (orange circle). Memory for the features was scaled between 0 (incorrect) and 1 (correct), with values in between graded by confidence (sound) or precision (color and scene) of the response, and summed to provide an overall memory quality score between 0 and 3 for each event.

### EEG Data Acquisition

EEG data were acquired using a 64-channel BioSemi ActiveTwo system. Electrodes were placed and secured on an elastic cap with conducting electrode gel in accordance with the International 10-20 system. Active electrodes were grounded against the common mode sense (CMS) and driven right leg (DRL) electrodes. Electrodes were additionally referenced against two external electrodes placed on the mastoids. EEG signals were recorded with a sampling rate of 512 Hz and with a recording bandpass filter from DC to 104 Hz. Additionally, electrodes were placed below and lateral to the eyes to detect eye movements and blinks using horizontal and vertical electrooculograms (EOG).

### EEG Data Processing

EEG data were preprocessed using the EEGLAB toolbox (Delorme & Makeig, 2004) in MATLAB. Data were resampled at 512 Hz with a high pass filter of 1 Hz. Bad channels, including those that flatlined, had excessive noise, or drifted, were identified and removed. Specifically, channels that had a flatline period greater than 5 seconds were identified as abnormal and removed. If a channel had greater than 4 standard deviations of line noise relative to its signal, based on all channels, it was removed. Retrieval data were baseline-corrected (on the basis of the 500-ms pre-stimulus interval), segmented into trial epochs from -1 to 6 seconds to facilitate wavelet-based time-frequency decomposition of lower frequencies (Cohen, 2014), and time locked to stimulus onset (the item shown in greyscale during the 4 second covert recall period, see Figure 1). Next, an independent component analysis (ICA) was used to automatically label additional noise components (e.g., muscle artifacts, heart beat, eye blinks) with at least 90% confidence (Pion-Tonachini et al., 2019), which were manually inspected and removed from the data. Following ICA, data were then interpolated across the removed bad channels, and the resulting data were run through the automatic data rejection algorithm once more to remove epochs that were greater than 2 standard deviations from the mean across all channels or greater than 6 standard deviations from the mean across a single channel (Delorme et al., 2007). Data were then visually inspected to exclude any remaining artifacts. Following preprocessing, one participant still had 11 noisy channels that were not identified by the automatic channel removal and was excluded from further analyses. Six additional participants were excluded due to noisy EEG data and because they had fewer than 70% of retrieval trials remaining after preprocessing. Overall, 23 participants were included in all analyses (see Participants, above).

### Behavioral Analyses

In our behavioral analyses, we first considered how well participants could remember each individual feature associated with an event - the sound, color, and scene location. Memory for the sound was coded as 0 if the participant’s response was incorrect, 0.5 if their response was correct but low confidence, and 1 if their response was correct and high confidence. For the color and scene features, responses were initially coded based on error (-180 to 180 degrees) between the participant’s response and the correct value on the color wheel or within the scene panorama (i.e., remembered feature value – encoded feature value; see Figure 1C). We then applied a mixture model of a von Mises distribution and uniform distribution to response errors across all subjects, separately for color and scene features, consistent with our prior work (Cooper & Ritchey, 2019). Using this model, we defined two threshold values for memory success where a response was less than 50% likely to be a guess (defined as fitting the uniform distribution). A correct response for color was defined as being within ±48 degrees of the correct location on the color wheel. A correct response for scene was defined as being within ±30 degrees of the correct location in the 360-degree panoramic image. Finally, responses were scaled to be between 0 and 1, where a response was coded as 0 if it was “incorrect” (at least a 50% chance of it being a guess) and between 0 and 1 if it was “correct” (less than a 50% chance of it being a guess), weighted by the error, such that scores closer to 1 reflect a response closer to the correct value (more precise). We then assigned each trial a composite “memory quality” score from 0 to 3, which was the summed score of the sound, color, and scene features (see Figure 1C).

In line with dependency analyses reported in Cooper and Ritchey (2019), we also sought to investigate how features were bound in memory, both in terms of binary success and precision. To this end, we first calculated the trial-to-trial dependency of binary memory success (i.e., correct vs. incorrect) for each pair of features (i.e., color-scene, color-emotion, and scene-emotion). This dependency score reflects the proportion of trials in which features were recalled or forgotten together within each participant (P_AB_ + P_A’B’_), which was corrected by the level of dependency predicted by an independent model accounting for overall memory accuracy, where better memory would lead to stronger correlations between the feature pairs (P_AB_ + P_A’_P_B’_; Bisby et al., 2018; Cooper & Ritchey, 2019; Horner & Burgess, 2013, 2014). We then calculated within-participant Pearson’s correlations (Fisher *z* transformed) between the precision of correct color and scene memory and memory success, defining trial-specific precision as the reversed absolute error of correct (i.e., memory success) trials such that higher values reflect greater precision. To test the dependence of color and scene precision, we restricted trials to those in which both features were successfully recalled. Finally, we also calculated within-participant Pearson’s correlations (Fisher *z* transformed) between each pair of features on the scaled memory scores that together constitute the composite “memory quality” score. Dependency scores for each feature pair were compared against *mu* = 0 in a series of one-sample t-tests.

### EEG Analyses

EEG analyses tested how continuous memory measures parametrically modulated event-related potentials and oscillatory waveforms. Trials were scored as described above (see Behavioral Analyses). For both event-related potentials (ERP) and oscillatory activity, we conducted analyses for the composite memory quality scores first and then for each memory feature (i.e., color, scene, and emotion) separately.

We first conducted a stimulus-locked analysis of ERP data. We selected an *a priori* region of interest (ROI: CP5, CP3, CPz, CP4, CP6, P3, Pz, P4, CP1, CP2, P1, P2) between 500-800 ms following stimulus onset, the time window of the late positive component (LPC; (Addante et al., 2012; Leynes & Crawford, 2018; Murray et al., 2015; Walsh et al., 2016). Using the LIMO EEG toolbox (Pernet et al., 2011), we ran two continuous Ordinary Least Squares (OLS) regressions per participant to see if there was a relationship between ERP power and memory performance. These two regression models were specified as follows: *(i)* overall memory quality, whereby ERP power was predicted by the composite memory score (*Y* = β_0_ + β_1--_*X*_1_ + L); and *(ii)* color, scene, and emotion, whereby ERP power was predicted by memory for each event feature (*Y* = β_0_ + β_1--_*X*_1_ + β_2_*X*_2_ + β_3--_*X*_3_ + L). Resulting beta coefficients from the overall memory quality model were first averaged across channels in our posterior ROI to test the relationship between ERP power and memory performance from stimulus onset to 1000 ms post-stimulus, running a one-sample *t*-test at each time point controlling the false discovery rate.

We then averaged ERP model coefficients from channels within our posterior ROI and across all time points in our 500 – 800 ms LPC window of interest, resulting in time-averaged *beta* values per subject for the composite memory quality score, as well as for each feature in memory separately. For memory quality, we identified one extreme outlier (±1.5 * interquartile range) and found that the distribution of data departed significantly from normality (*W* = 0.87, *p* < .01), so we opted to conduct a nonparametric one-sample Wilcoxon signed rank test to compare the averaged ERP model coefficients against *mu* = 0. The spatial distribution of this memory quality effect within the LPC window was then visualized using topographical mapping. Importantly, to better elucidate the spatial distribution of recollective experience compared to all-or-nothing remembering, we also visualized the memory quality effect within the LPC window for averaged forgotten trials (i.e., memory quality score = 0; “incorrect”) compared to averaged trials of all levels of memory quality (i.e., memory quality score > 0) using topographical mapping.

Finally, for each of the three memory features (i.e., scene, color, and emotion), we conducted one-sample comparisons on averaged ERP model coefficients within the LPC window for our posterior ROI against *mu* = 0. Note that the distribution of data for scene memory departed significantly from normality (*W* = 0.90, *p* < .05) and we identified one extreme outlier so we opted to conduct a nonparametric one-sample Wilcoxon signed rank tests for all memory features. We also conducted pairwise comparisons between features for ERP model coefficients. The spatial distribution of memory feature effects were also visualized using topographical mapping. For all topographical maps of ERP results, significant channels were determined using one-sample *t*-tests and indicated in figures with white crosses (α = 0.05).

We additionally conducted a time-frequency analysis focusing on a frequency range from 5 to 30 Hz, spanning the theta (5 - 7 Hz), alpha (8 - 12 Hz), and beta (13 - 30 Hz) frequency bands (Buzsáki, 2006). Frequency decomposition from the EEG data was obtained for each condition, using Morlet wavelets (Cohen, 2014) and a fixed epoch length (i.e., 500 ms pre-stimulus onset to 4000 ms post-stimulus onset, comprising the covert recall period). The sliding window length varied by frequency, from 5 cycles at 5 Hz (1000 ms window) to 6 cycles at 30 Hz (200 ms window). Using the LIMO EEG toolbox, we again ran two continuous OLS regressions per participant for *(i)* memory quality (*Y* = β_0_ + β_1--_*X*_1_ + L) and *(ii)* color, scene, and emotion features (*Y* = β_0_ + β_1--_*X*_1_ + β_2_*X*_2_ + β_3--_*X*_3_ + L) to determine if a relationship between time-frequency representations of power and memory performance exists. The resulting model coefficients were then run through a series of one-sample *t*-tests. Specifically, we conducted a permutation test by electrode at each frequency and time and then used cluster correction to identify where the relationship between time-frequency power and memory performance reliably differed from 0. Since we were interested in positive and negative clusters separately, we set a threshold of α = .025 (two-tailed) to determine significant clusters. This correction method accounts for spatial and temporal dependencies in the data by identifying clusters of contiguous significant values and correcting for multiple comparisons within these clusters. Bootstrapping with resampling was employed to estimate the null distribution for the cluster-based correction. Finally, we also sought to investigate the relationship between significant negative clusters identified for overall memory quality and clusters identified for each feature in memory, to better understand how time-frequency power related to retrieval of color, scene, and emotion features contributed to alpha/beta desynchronization related to overall memory performance. To this end, we determined whether significant cluster-corrected alpha/beta desynchronization effects for each feature overlapped in time and frequency with negative clusters identified for overall memory quality.

In addition to identifying the neural markers of quality and content in memory recollection using ERPs and time-frequency analysis, we also sought to delineate the relationship between these markers. To date, there is limited research on whether a relationship exists between ERPs and time-frequency responses in the context of memory recollection, though some findings suggest a correspondence in memory retrieval more broadly (see, for e.g., Düzel et al., 2005 in the context of recognition memory with magnetoencephalography). First, we calculated the average effect size for memory quality within the LPC window (i.e., the time window of interest in our ERP analyses described above) for each channel, averaging across participants. Second, for each channel, we similarly calculated the average effect size within time-frequency windows of significant clusters of alpha/beta desynchronization for memory quality, as identified in the time-frequency analyses. Next, we ran a simple linear regression in which we sought to determine whether the average effect within each negative cluster in the time-frequency analysis predicted the average ERP effect during the LPC window (*Y* = β_0_ + β_1_*X*_1_ + β_2_*X*_2_ + β_3_*X*_3_). In other words, do channels that show a large desynchronization effect also show a large LPC effect? We predicted that if ERPs and time-frequency patterns are largely overlapping in the neural processes they measure, a significant relationship would be present. Otherwise, if no relationship is found, then it would suggest that these neural markers may be underlying different aspects of memory recollection.

## Results

### Behavior

Using our mixture model fit across group averaged responses, we first determined how many participant responses for each behavioral measure were within the threshold to be marked as “correct,” i.e., with less than a 50% probability of being from the uniform “guess” distribution. Across participants, 69.96% (*SE* = .029) of color responses were marked as correct, that is, within 48 degrees of the studied color (Figure 2A). Across participants, 69.79% (*SE* = .028) of scene responses were marked as correct, that is, within 30 degrees of the location of the item in the panoramic image (Figure 2B). Additionally, 75.86% (*SE* = .022) of sound responses were correct, that is, correctly identified as a negative or neutral sound during retrieval. We also examined the distribution of composite memory quality scores, which were skewed toward higher scores but were well-distributed across the range from 0 to 3 (Figure 2C). This indicated that participants were able to remember multiple features of the overall encoded event but with enough variability to capture modulation of ERP and oscillatory effects. All feature pairs showed significant memory dependency for binary retrieval success (Figure 2D), indicating that successful retrieval of one feature was likely to lead to the retrieval of another feature (*t*s(22) > 4.17, all *p*s < .001). We also found that retrieval success of scene features significantly benefited the precise recollection of color features (Figure 2E; *t*(22) = 4.91, *p* < .001). In contrast, we found that precision memory for color and scene features were unrelated (*t*(22) = 0.17, *p* = 0.87) and retrieval success of color features did not benefit the precise recollection of scene features (*t*(22) = 1.70, *p* = 0.10). Finally, we found that all feature pairs were significantly related when considering scaled memory scores, which reflect a combination of success and precision (Figure 2F; *t*s(22) > 6.44, all *p*s < .001).

**Figure 2.**
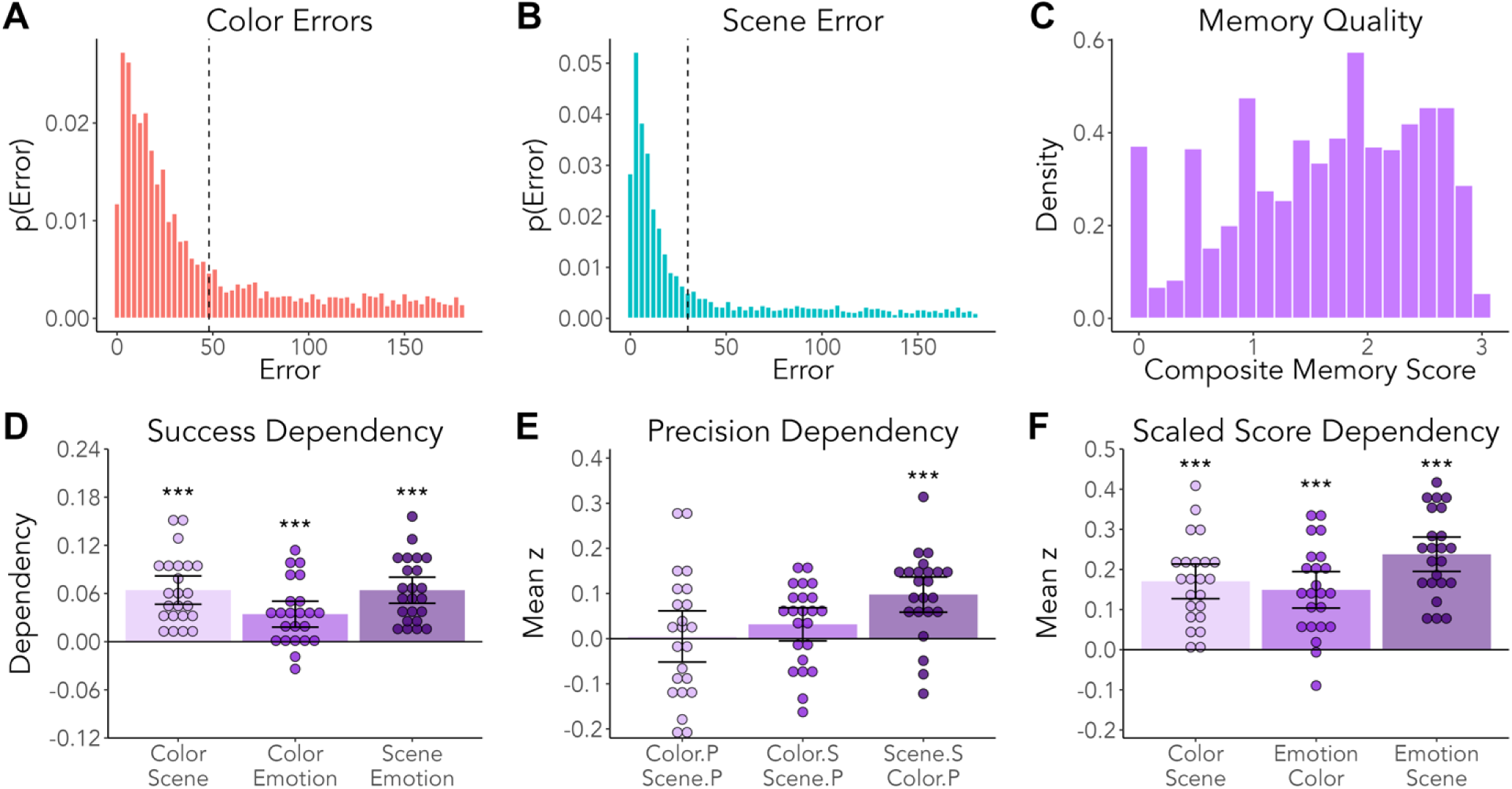
Aggregate absolute error distribution (response - target) for color and scene, and distribution of memory quality scores. (A) Color responses were marked as correct if the response was within 48 degrees of the correct location on the color wheel, as denoted with the dashed vertical line. (B) Scene responses were marked as correct if the response was within 30 degrees of the correct location in the panoramic image, as denoted with the dashed vertical line. (C) Distribution of composite memory quality scores. (D) Corrected memory dependency between feature pairs across trials within participants, in terms of memory success (i.e., correct vs. incorrect). (E) Mean Fisher z transformed Pearson’s correlations between the precision (P) of remembered color and scene trials and successful (S; correct vs. incorrect) retrieval of those features. (F) Mean Fisher z transformed Pearson’s correlations between scaled memory scores for each feature pair. (Error bars represent mean ±95% CI, ***p < .001)

### Event-Related Potentials

To investigate whether the late positive component (LPC) was modulated by multidimensional memory quality, we used a continuous regressor of ERP power modulated by overall memory quality. We ran a one-sample *t*-test at each time point from 0-1000ms to test if any individual time points in our posterior ROI were significantly different from 0 (Figure 3A), but no time points were significant after correction for multiple comparisons by controlling the false discovery rate. Averaging across the LPC time window (500 – 800 ms post-onset) and across channels in our posterior ROI, memory quality betas were positive and significantly differed from zero (*V* = 218, *p* = .014, *r* = .51, Wilcoxon’s sign-rank test) (Figure 3B). The removal of the extreme outlier value of one participant did not change the significant effect (*t*(21) = 2.34, *p* = .029, *d* = .50).

**Figure 3.**
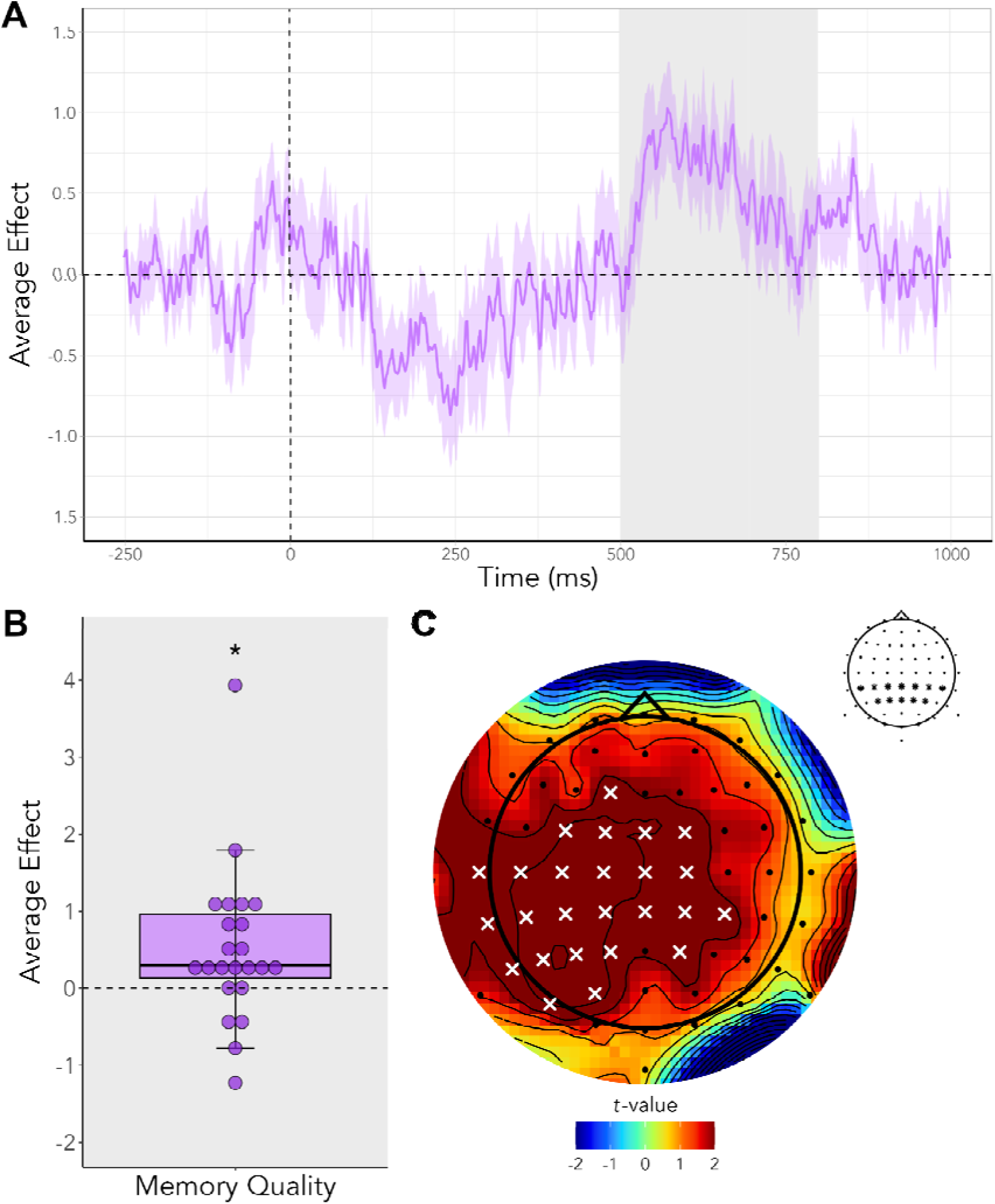
Group-averaged ERP waveform for memory quality in our posterior ROI. (A) The average effect of composite memory quality on ERP power plotted from -250 ms pre-onset to 1000 ms post-onset for all channels. The time window of interest for the LPC (500 – 800 ms following stimulus onset) is highlighted in gray. Shaded areas represent the average effect of composite memory quality on ERP power ± SEM. (B) The average effect of composite memory quality on ERP power averaged acros timepoints for channels in the posterior ROI within the LPC time window (i.e., 500– 800 ms post-onset). Averaged memory quality ERP power in this time window is significantly different from 0 (error bars represent ± SEM, *p < .05). (C) Topographical mapping of LPC effects for memory quality across all channels within the LPC time window, with significant channels (α = 0.05) indicated by white crosses. Channels in our ROI are indicated with asterisks in the topographical mapping legend in the upper right corner.

To complement the above ROI analysis and to verify the spatial distribution of LPC effects for memory quality, we additionally examined activity across all channels during the LPC time window (Figure 3C). Specifically, we averaged the beta values from the composite memory quality model within each channel across all time points from 500 – 800 ms. Activity was widely distributed across channels in our posterior ROI, in addition to select fronto-central channels. A series of one-sample *t*-tests (*mu* = 0) revealed significant activity in channels within our posterior ROI (all *p*s_uncorr_ < .05), except in CP6, P4, and Pz.

We next investigated how the LPC was modulated by memory precision for the individual item features (i.e., color and location within each scene), as well as by confidence for the emotional sound association. Our analyses here compared the LPC effect for channels in our posterior ROI to *mu* = 0, following up on the effect of overall memory quality found above during this time window (Figure 4A).

**Figure 4.**
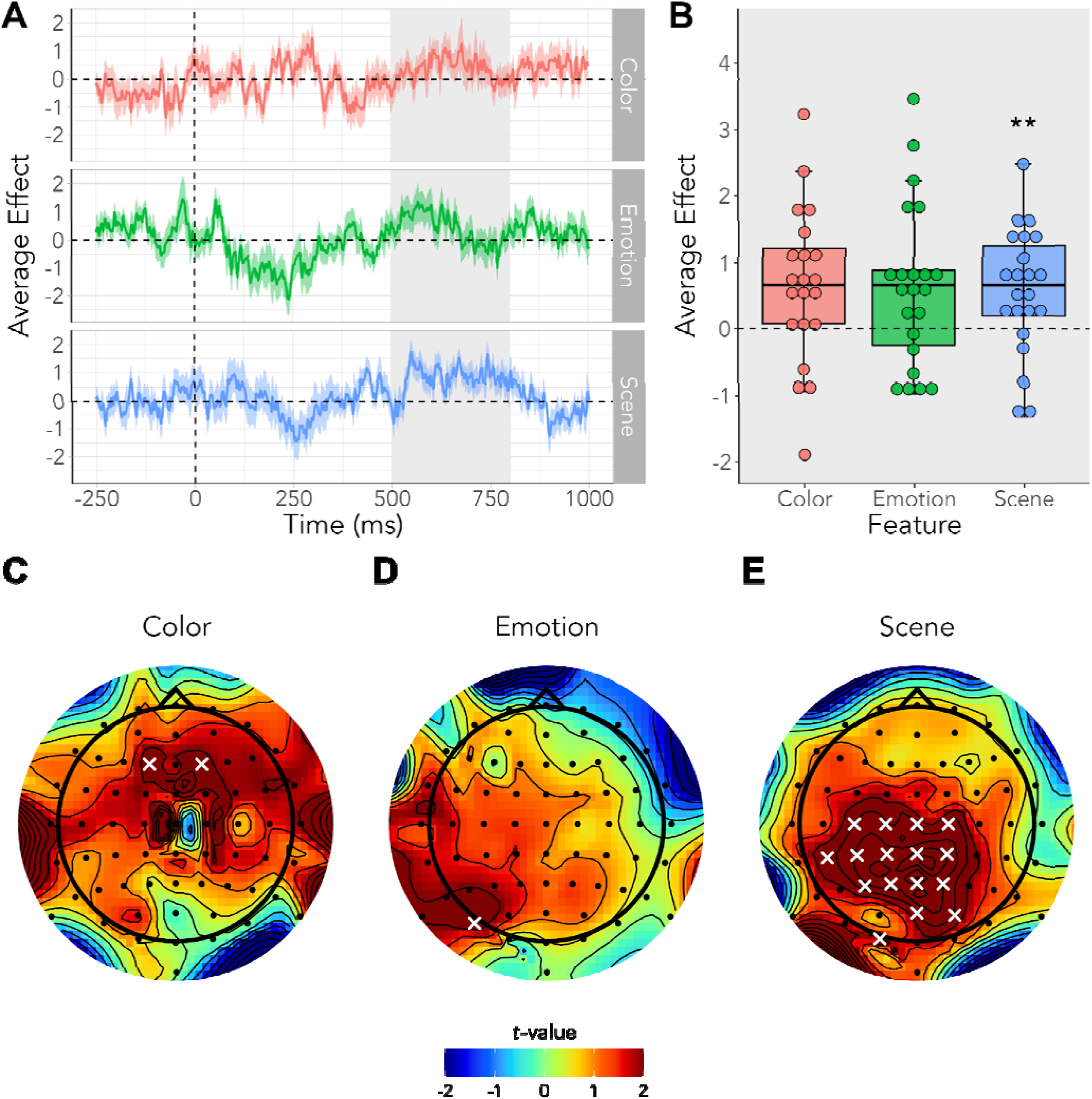
Group-averaged ERP waveforms for color, emotion, and scene in channels of our posterior ROI. (A) ERP power plotted from 500 ms to 800 ms post-onset (i.e., LPC window) for channels in the posterior ROI for each feature. Shaded areas represent averaged ERP power ± SEM. (B) ERP power averaged across timepoints for channels in the posterior ROI within the LPC time window for each feature (**p < .01). Topoplots for feature memory averaged within the LPC window, with (C) color, (D) emotional sound, and (E) scene features shown, respectively. Significant channels (α = 0.05) are indicated by white crosses.

There was no significant relationship between the ERP power and memory for color (*V* = 186, *p* = 0.15, Wilcoxon’s sign-rank test) or emotion (*V* = 184, *p* = 0.17, Wilcoxon’s sign-rank test). We did, however, find a significant, positive relationship between scene memory precision and ERP power (*V* = 229, *p* = .004, *r* = .58, Wilcoxon’s sign-rank test) (Figure 4B). Pairwise comparisons between features did not reveal any significant differences in ERP power within the LPC window in our posterior ROI (all *p*s > 0.05). Furthermore, to determine whether the LPC may have been modulated by memory precision for color or by confidence for the emotional sound association outside of our posterior ROI, we conducted a follow-up analysis against *mu* = 0 averaging across all channels. However, there was no significant relationship between the averaged ERP power and memory for color (*t*(22) = 0.42, *p* = .14), nor for emotional sound (*t*(22) = .24, *p* = .38), when averaging across all channels.

Echoing our topographical analyses for memory quality, we then examined ERP power across all channels for each of the features of memory precision (i.e., color, scene, and emotional sound). During the LPC window, we did not find significant activity in channels within our posterior ROI for color (Figure 4C), or for emotion (Figure 4D) memory. We found significant widespread activation in our posterior ROI during the LPC window for scene memory (all *p*s_uncorr_ < .05; Figure 4E). This suggests that memory for scenes might drive the modulation of the LPC window in our posterior ROI (i.e., left parietal regions).

### Oscillatory Activity

#### Memory Quality

We first examined the relationship between overall memory quality and oscillatory activity across all channels, all time points between 0 ms – 4000 ms (i.e., the covert recall period), and all frequencies between 5 – 30 Hz (see Figure 5A). Positive and negative clusters reflecting alpha/beta desynchronization are reported in Table 1 below. We primarily focused our analyses on significant negative clusters spanning alpha and beta frequencies across the 4000 ms recall period, which were especially evident in central-posterior channels. Specifically, results showed a significant negative effect surviving cluster-based correction for multiple comparisons (Maris & Oostenveld, 2007) early in the recall period, at 798 – 918 ms and between 15.74 – 30.00 Hz. The relationship between overall memory quality and desynchronization in the recall period was also evident in a significant negative cluster between 1125 ms – 2277 ms across alpha and beta frequency ranges (9.53 – 30.00 Hz). Finally, we found a significant negative cluster late in the recall period (2840 – 3105 ms) broadly spanning alpha and beta frequencies as well as the theta frequency range (5.27 – 16.91 Hz) (see Figure 5B). Two positive clusters spanned a low alpha frequency range, early in the covert recall period.

**Figure 5.**
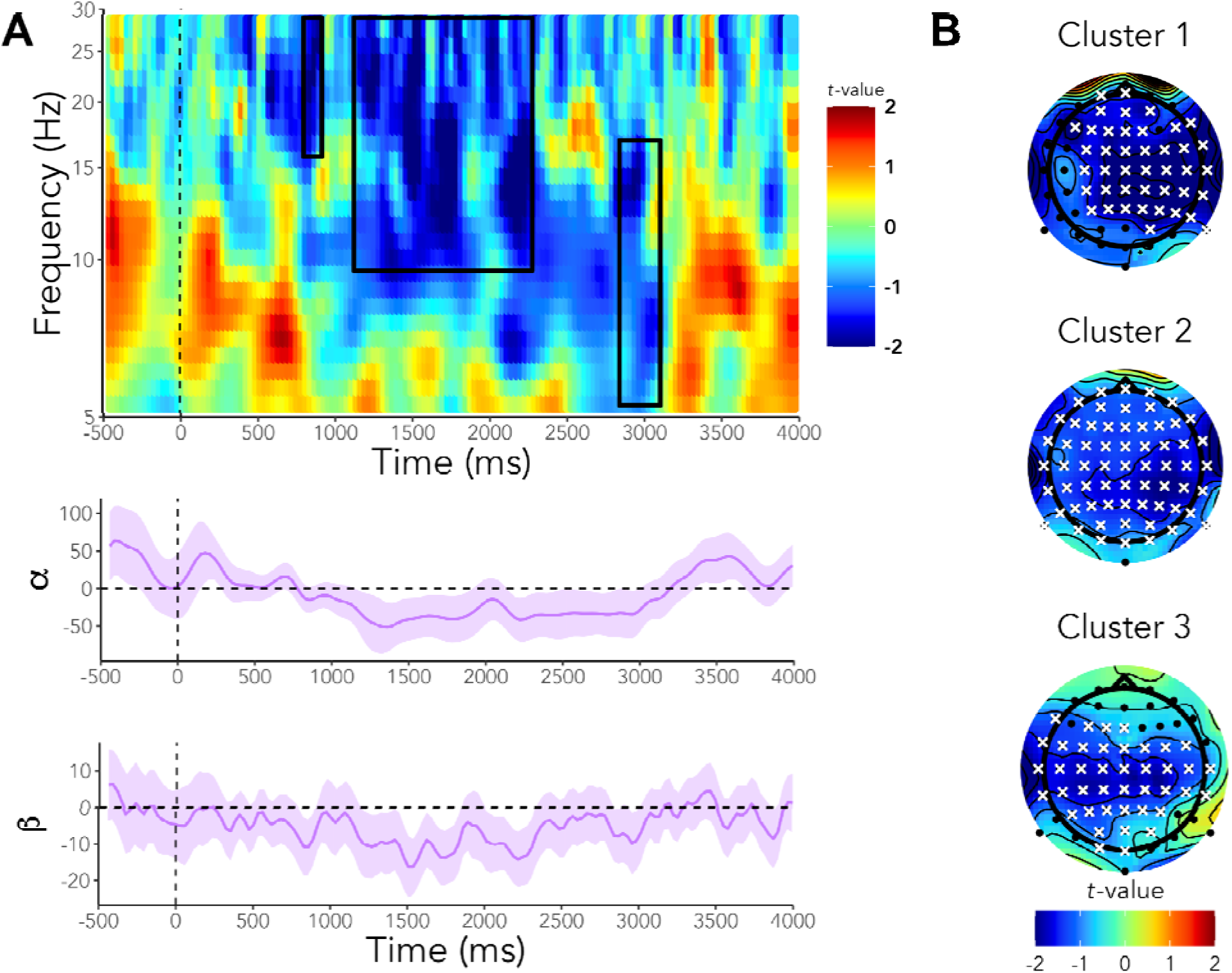
The relationship between overall memory quality and alpha/beta desynchronization during the covert recall period (i.e., 0 – 4000 ms). (A) Time-frequency representation for the covert recall period spanning frequencies between 5 – 30 Hz (log transformed to improve visualization of lower frequencies). Significant clusters are highlighted by the solid-line boxes. Alpha (α) and beta (β) frequencies are plotted in the lower panel across the covert recall period as a function of their relationship with memory quality below the time-frequency representation. (B) Topoplots representing three significant negative clusters spanning alpha and beta frequency ranges between 798 – 918 ms, 1125 – 2277 ms, and 2840 – 3105 ms, respectively.

**Table 1.** Significant positive and negative effects surviving cluster-based correction for multiple comparisons during the covert recall period.

| Cluster | Frequency (Hz) | Times (ms) | Channels |
| --- | --- | --- | --- |
| **Positive** |  |  |  |
| 1 | 7.15 - 9.53 | 177 - 297 | C1, CP1, CP2, CPz, Cz, P1, P2, Pz, P4, PO4, POz, CP3, CP5, P3, P5 |
| 2 | 5.37 - 8.87 | 592 - 838 | AF4, AF8, F6, F8, FT8, AF3, AFz, C6, CP5, F1, F2, F3, F4, FC4, FC6, Fp2, Fz, P3, P5, T8, C5, F5, F7, Fpz, FT7, PO3, T7, TP7, FC5, O2, PO4, PO8, POz, Iz, O1, Oz, PO7 |
| **Negative** |  |  |  |
| 1 | 15.74 - 30.00 | 798 - 918 | CP6, P6, T8, TP8, CP4, P10, P2, P4, AF3, AF4, AFz, C1, C2, C4, C6, CP1, CP2, CP3, CPz, Cz, F1, F2, F3, F5, FC1, FC2, FC3, FC6, FCz, FT8, Fz, P1, P8, PO4, Pz, C3, FC4, Fp1, Fpz, P3, F6, F8 |
| 2 | 9.53 - 30.00 | 1125 - 2277 | Cz, F1, F2, F4, FC1, FC2, FC4, FCz, Fz, C1, C2, C4, FC6, CP2, CP4, F6, FT8, T8, CP6, F3, FC3, P2, P4, C6, CP3, O2, P1, P3, P6, PO4, PO8, POz, TP8, C3, CP1, F8, P5, P8, CP5, CPz, Pz, T7, TP7, PO3, F5, AF3, AF4, AF7, AF8, C5, F7, FC5, Fp1, Fp2, FT7, O1, Oz, P7, P10, P9, PO7, AFz, Fpz |
| 3 | 5.27 – 16.91 | 2840 - 3105 | C1, C2, C3, C4, C5, C6, CP1, CP2, CP3, CP4, CPz, Cz, F7, FC3, FC4, FC5, FC6, FT7, T8, CP5, FC1, FC2, T7, TP7, FCz, P1, P3, PO3, F1, F3, Fz, O1, Oz, P5, POz, CP6, Pz, TP8, P4, P2, PO4 |

#### Episodic Features

The subsequent analyses focused on whether changes in low-frequency power (i.e., alpha and beta ranges) were modulated by the content and precision of recollection during the covert recall period (0 - 4000 ms). To this end, we initially examined the relationship between features in memory and oscillatory activity across all channels, all time points between 0 ms – 4000 ms (i.e., the covert recall period), and all frequencies between 5 – 30 Hz. A series of significant negative clusters consistent with evidence for alpha/beta desynchronization were identified for each memory feature across the covert recall period (see Figure 6A).

**Figure 6.**
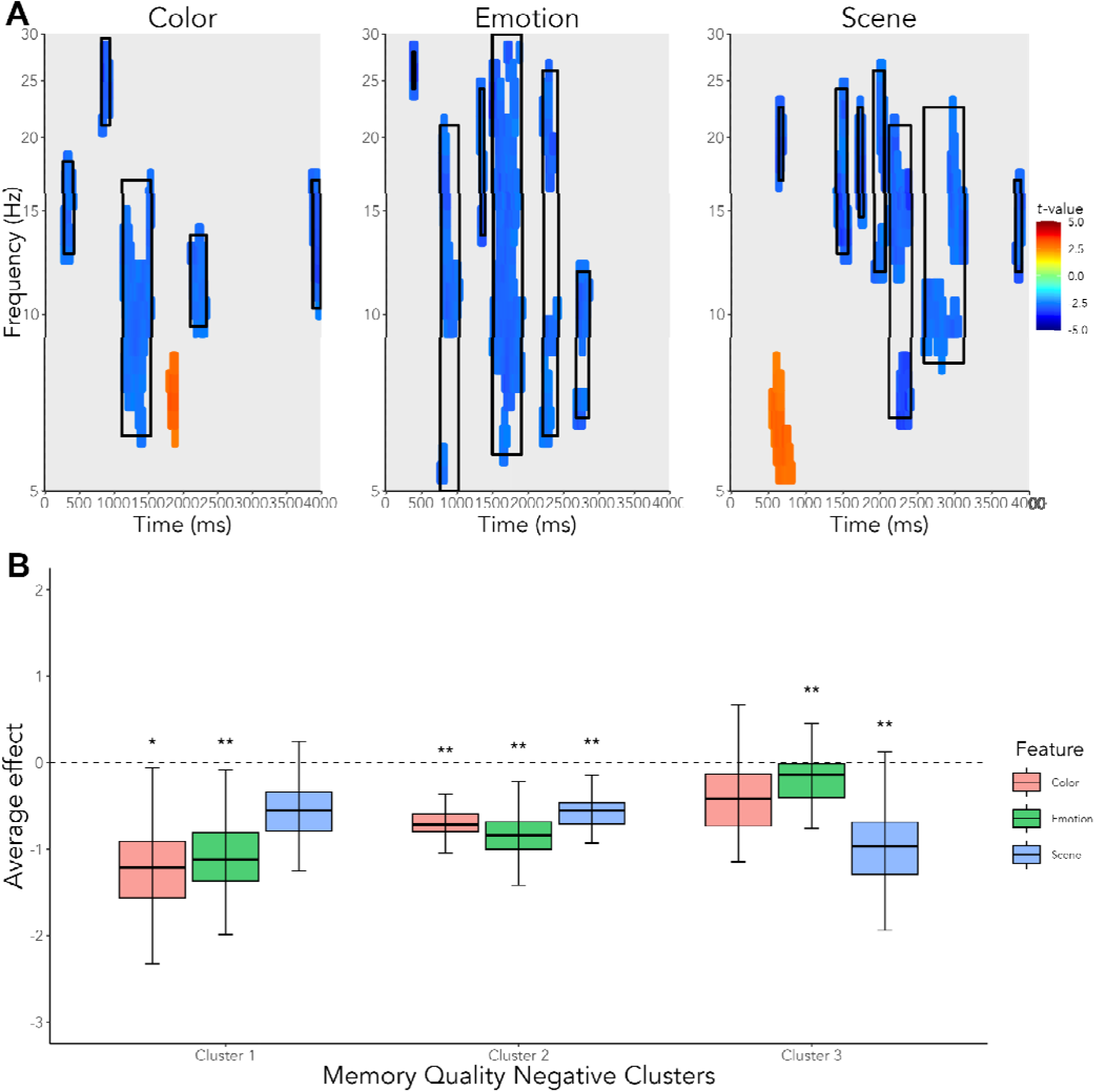
The distribution of average negative modulatory memory effects on alpha and beta frequency bands, shown for color, emotion, and scene features. (A) Significant negative clusters indicating alpha/beta desynchronization for color, emotion, and scene, respectively. Negative clusters are indicated by boxes in each figure. (B) Average effect surviving cluster-based correction for color, emotion, and scene features within significant overall memory quality negative clusters. (*p < .05, **p < .01).

We then looked at the correspondence between the three negative clusters identified above for overall memory quality and negative cluster effects for each of the individual features. In this way, we could determine how the precise retrieval of memory features underlies alpha/beta desynchronization patterns consistent with overall memory quality (see Figure 6B). For the first cluster (i.e., 799 – 918 ms; 15 – 30 Hz) we found a corresponding negative effect surviving cluster-based correction for multiple comparisons for color and emotion, but not scene memory. Cluster 2 (i.e., 1125 ms – 2277 ms; 9.53 – 30 Hz) corresponded to negative effects for all features, suggesting that alpha/beta desynchronization during this middle time window of the covert recall period is driven by precise retrieval of all features in memory. Finally, for cluster 3 (i.e., 2840 – 3105 ms; 5.27 – 16.91 Hz) we found a corresponding negative effect primarily for scene memory, but not for color memory, suggesting a shift in alpha/beta desynchronization reflecting retrieval of color features from memory in the early stages of the covert recall period towards retrieval of scene features from memory in the later stages of the covert recall period.

### Event-Related and Time-Frequency Correlation Analyses

Because we found evidence for overall memory quality modulating the LPC in stimulus-locked analysis of ERP data and low-frequency bands in our time frequency analysis, we finally sought to investigate whether these two sets of neural markers were related to one another. To this end, we ran a linear regression in which, across channels, averaged beta coefficients for each of the significant negative alpha/beta clusters for memory quality within the covert recall period predicted the averaged beta coefficients in the LPC window. This tested whether the LPC and desynchronization effects were colocalized, with channels showing a large alpha/beta desynchronization effect also showing a large LPC effect. Indeed, there was a significant relationship between the effects of memory on alpha/beta desynchronization and its effects on the LPC (*F*_3,60_ = 40.53, *p* < .001, *R^2^*= .670). Examining the individual predictors indicated that cluster 1 (*t* = -1.53, *p* = 0.13) and cluster 2 (*t* = -1.15, *p* = 0.26) were not significant predictors, but cluster 3 (i.e., 2840 – 3105 ms; 5.27 – 16.91 Hz) was a significant predictor of averaged beta coefficients in the LPC window (*t* = -9.83, *p* < .001, see Figure 7).

**Figure 7.**
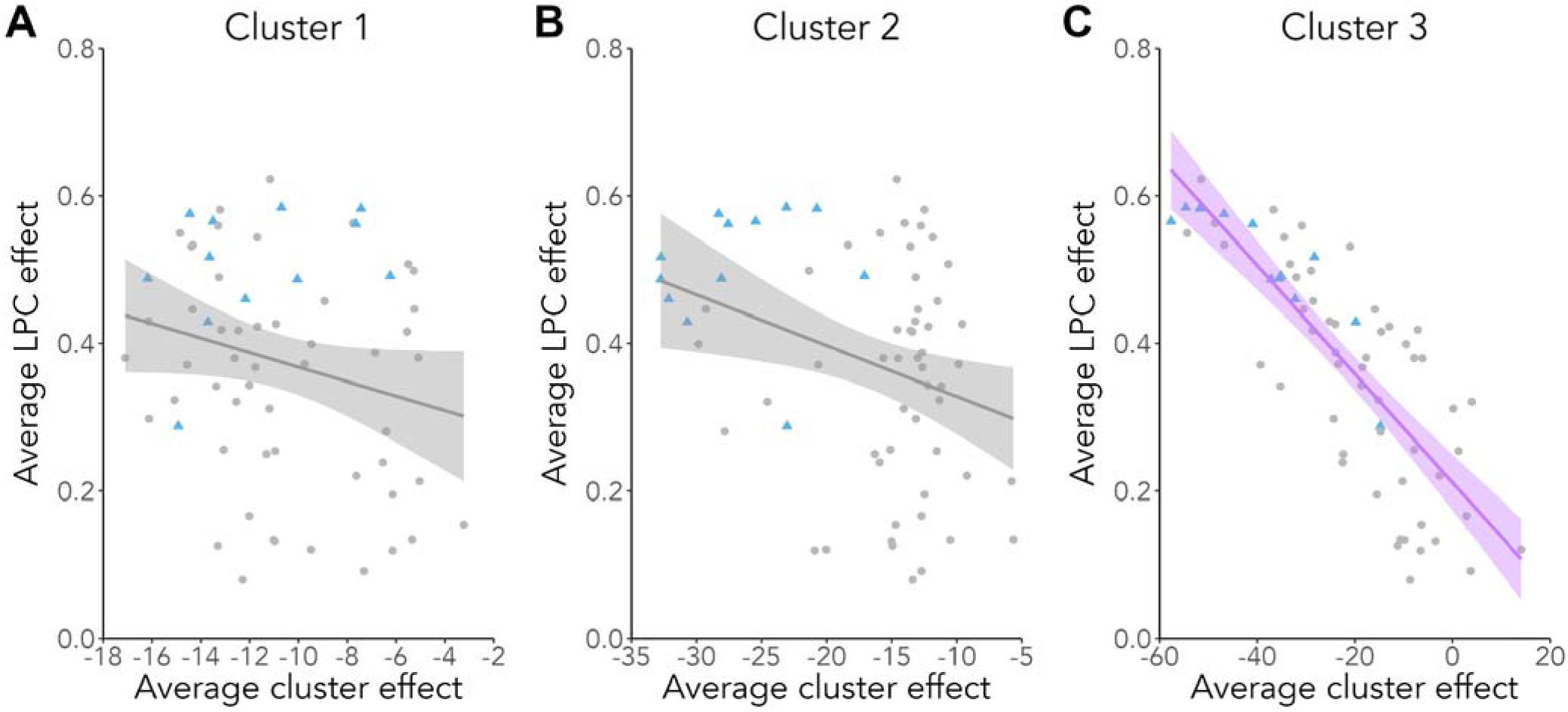
The relationship between averaged beta coefficients for each channel within the LPC window and cluster 3 indicating significant alpha/beta desynchronization during the covert recall period. (A-B) No significant relationship between the LPC effect and cluster 1 or cluster 2 were observed. (C) Channels which showed strong alpha/beta desynchronization in cluster 3 reflected more activity modulated by memory quality during the LPC window. For A-C, each point represents a unique channel, with blue triangles representing channels within our posterior ROI.

## Discussion

In the current EEG study, we investigated how two EEG signatures of recollection – the LPC and alpha/beta desynchronization – were modulated by the multidimensional quality of episodic memory. We identified a relationship between overall memory quality and ERP activity within the LPC window, between 500 and 800 ms post-stimulus onset, predominantly in the left posterior region of the brain. Importantly, left posterior electrodes within the LPC window were also sensitive to the precision of scene memory, but not to the precision of memory for color or emotion features. Complementing our ERP findings, analyses of oscillatory activity during the covert recall period revealed evidence of three distinct clusters reflecting alpha/beta desynchronization modulated by memory quality. Moreover, we identified specific clusters of alpha/beta desynchronization modulated by memory precision for specific features (i.e., color, emotion, and scene) and demonstrated how these clusters might contribute to patterns of alpha/beta desynchronization related to overall memory quality. Finally, we found that alpha/beta desynchronization modulated by memory quality late in the covert recall period predicts ERP activity modulated by overall memory quality within the LPC window. These findings are discussed in depth below.

Our findings of ERP activity modulated by overall recollection quality support the idea that the LPC effect in left posterior electrodes – corresponding to parietal regions of the brain – is not limited to all-or-nothing, old/new memory. The specific pattern of our results extends findings from prior ERP studies of the LPC window beyond binary measures of retrieval (see e.g., Curran & Cleary, 2003; Vilberg & Rugg, 2009) and towards precision-based continuous measures of multi-feature recollection. Importantly, our multi-feature paradigm also allowed us to investigate the content of bound memory representations, beyond single feature associations. Behaviorally, we found that these bound memory representations seem to contain gist-like information dependent across features, but that the specific resolution of each feature appears to be independent. Our ERP results also largely converge with prior fMRI studies showing functional engagement of parietal regions with memory recollection, including angular gyrus (Bellana et al., 2023; Ramanan et al., 2018; Rugg & Vilberg, 2013) and precuneus (Hebscher et al., 2018; Sreekumar et al., 2018). More specifically, recall-related patterns of fMRI activity in the lateral parietal cortex differentiate between individual events and have been shown to be sensitive to the subjective experience of vividness in memory (Kuhl & Chun, 2014). Here, our ERP results provide further insight into the temporal dynamics of recollection, demonstrating a distinct pattern of parietal modulation specific to the LPC window (i.e., beginning at 500 ms post-stimulus onset). More broadly, the posterior medial episodic network – a subsystem of the default network (Ritchey & Cooper, 2020) – is consistently coactivated with the hippocampus during recollection (Benoit & Schacter, 2015; Rugg & Vilberg, 2013; Spreng et al., 2009) and comprises several parietal regions. Recently, we reported that hippocampal connectivity with bilateral parietal cortex was specifically modulated by memory quality, while participants completed the same multi-feature paradigm reported in the current study (Cooper & Ritchey, 2019). Consistent with these findings, intracranial EEG recordings in human epilepsy patients suggest that sensory cues proceed towards the hippocampus between 0 and 500 ms post-stimulus onset, enabling cortical memory reinstatement to unfold subsequently (Staresina et al., 2012; for review see Staresina & Wimber, 2019). Here, we show that this memory reinstatement likely involves recalling high-dimensional representations at varying levels of retrieval quality. In future studies, combining EEG approaches with network connectivity analyses of fMRI data using this multi-feature paradigm could further elucidate the relationship between hippocampal-dependent episodic recollection and modulation of the LPC in parietal regions.

In addition to finding a relationship between overall memory quality and modulation of the LPC, we demonstrated that memory precision for specific episodic features also modulated the LPC. In particular, scene location memory in our study seemed to drive modulation of posterior and central regions, consistent with an earlier study from Murray et al. (2015). It is possible that because scene locations were discriminated in memory with higher precision, it may be a more sensitive measure of memory quality. However, we argue that this finding is not necessarily explained by task difficulty or, rather, the facility with which scene location memory was assessed compared to color memory. Although the thresholds for color and scene memory accuracy in our mixture model differed, within those thresholds we found comparable behavioral memory performance (i.e., roughly 70% correct) for scene and color memory. In other words, we only considered a subset of responses that we would expect were less than 50% likely to be guesses. Instead, perhaps a more plausible interpretation of this finding is that spatial context provides a scaffold for the unfolding of other episodic features during recollection (Robin, 2018; Robin et al., 2016) and that at the gist-level, the memory trace consists of bound information about distinct features which facilitate retrieval in a dependent manner. Indeed, our behavioral analyses indicate that successful retrieval of any feature in memory benefitted retrieval of other features in memory at the gist-level, but also that gist-level scene memory benefitted precision memory for color. Alternatively, our LPC results here could be explained by the location of electrodes within our posterior ROI (i.e., left parietal regions). Representations of spatial features may be particularly salient in the parietal cortex (see e.g., Husain & Nachev, 2007; Kesner, 2009), where we found the strongest LPC modulatory effects for scene location memory.

Our time-frequency analyses focused on a four-second covert recall period during which participants were instructed to bring to mind the spatial location, color, and emotional sound associated with an item previously encoded in the study. We found three clusters of activity reflecting alpha/beta desynchronization spanning an early, middle, and late period of the recall period modulated by overall memory quality. Existing frameworks suggest a temporal progression of early hippocampal theta/gamma synchronization to cortical posterior alpha/beta desynchronization as reflective of memory access facilitating memory elaboration and representation, respectively (Griffiths et al., 2019; Hanslmayr et al., 2016; Parish et al., 2018; Staresina & Wimber, 2019). Indeed, our time-frequency findings are largely consistent with prior studies of episodic memory retrieval and low-frequency desynchronization (Hanslmayr et al., 2012, 2016; Karlsson et al., 2020), and explicitly demonstrate that desynchronization scales with the quality of memory recollection. Moreover, we found that low-frequency desynchronization was also modulated by the content of memory recollection, with patterns of desynchronization observed for all features. Rapid decreases in alpha/beta power track with the reactivation of sensory information in memory (Waldhauser et al., 2016) and may serve to facilitate the flow of information about mnemonic content to relevant neural assemblies (Hanslmayr et al., 2012; Parish et al., 2018), potentially by reinstating similar patterns of desynchronization observed at encoding (Sutterer et al., 2019). Other work has proposed that alpha desynchronization may play a role in inhibiting global cortical activity, to enable specific activation of the regions associated with a target object during retrieval (Chiang et al., 2016; Klimesch, 2012). To better understand how alpha/beta desynchronization effects corresponded between overall memory quality and each episodic feature, we also considered the average cluster-corrected effects for features within the three identified memory quality clusters. In this way, we could determine how the precise retrieval of memory features underlies alpha/beta desynchronization patterns consistent with overall memory quality. We found a shift in alpha/beta desynchronization reflecting retrieval of color features from memory in the early stages of the covert recall period towards retrieval of scene features from memory in the later stages of the covert recall period. More specifically, the earliest cluster of alpha/beta desynchronization for memory quality corresponded with cluster-corrected effects for color and emotion, but not scene memory, whereas the second cluster for memory quality captured cluster-corrected effects for all three features in memory.

The final cluster – emerging late in the covert recall period – corresponded to scene and emotion memory, but not color memory. In the current experimental task, alpha/beta desynchronization early in the covert recall period may have reflected color because of the relevance of this feature to the presented cue (i.e., intra-item association with the grayscale item). Prior work suggests that attention can modulate the temporal dynamics of specific object features at encoding and retrieval (Mirjalili et al., 2021) such that participants in our study may have attended to intra-item features during retrieval first, followed by an unfolding of the spatial context. To our knowledge, our time-frequency findings are the first to demonstrate that alpha/beta desynchronization may reflect the dynamics of memory recollection for bound multidimensional representations and also for individual features.

A unique feature of the current study was that we examined ERPs and oscillatory patterns in the same dataset. Both the LPC (ERP) and alpha/beta desynchronization (time-frequency) have been reliably associated with recollection, but it is not fully understood in the literature to what extent these measures represent convergent markers of precision and content in memory. We found a relationship between memory-modulated activity in the LPC window and alpha/beta desynchronization during the late stages of the covert recall period. Specifically, channels demonstrating a strong desynchronization effect late in the covert recall period (2840–3105 ms within the covert recall period) also reflected more memory-modulated activity during the LPC window (500 – 800 ms). This finding suggests that both neural markers measure similar neural processes (Schneider & Maguire, 2018) and may reflect overlapping aspects of memory recollection. This is consistent with the observation that both of these markers appeared to be most sensitive to the precision of scene information, while broadly supporting the retrieval of multidimensional bound representations following stimulus onset. In related work, Chen and Caplan (2017) demonstrated that decreases in alpha-band oscillations during memory encoding of words predicted increases in LPC activity. However, inconsistent with our findings, the authors did not find such a relationship between LPC activity and decreases in alpha-bands oscillations during retrieval. It is possible that recognition memory for words may not sufficiently modulate low-frequency desynchronization to the same extent as multidimensional memory features, as the authors did not find a relationship between durations of alpha oscillations and memory performance across participants. Future research will need to more explicitly investigate the relationship between the LPC and alpha/beta frequency activity with the goal of disentangling how recollection is reflected in the temporal dynamics of cortical activity. In particular, spectral parameterization (e.g., FOOOF algorithm, Donoghue et al., 2020) may help to clarify whether the pattern of neural oscillation late in the covert recall period is independent from aperiodic, non-oscillatory shifts in activity. Some emerging evidence suggest that both oscillatory and aperiodic activity may be co-modulated during learning (Cross et al., 2022) and more broadly across different cognitive demands (Frelih et al., 2024). As such, we hope that our findings serve as a foundation for future research exploring the relationship between ERPs, aperiodic activity, and oscillations associated with memory retrieval.

## Conclusion

The present study revealed how two EEG markers of recollection are modulated by the content and quality of memory, further extending our understanding about how features are bound in episodic memory. Future research would benefit from examining individual differences in EEG markers of episodic recollection beyond old/new recognition, as past work has suggested some degree of between-subject variability in these neural markers (Dimsdale-Zucker et al., 2022; Murray et al., 2015) that could scale with the quality and content of recollection at an individual-level. Here, we demonstrate for the first time that these EEG markers extend to continuous, multidimensional measures of memory, capturing recollection as a representationally complex experience that integrates multiple event features. Altogether, our results show how these two EEG signatures of recollection dynamically capture the quality and content of precise, multidimensional memory recollection.

## Author contributions

**NLW:** Conceptualization, Methodology, Formal Analysis, Writing - Review & Editing, Visualization. **HS:** Conceptualization, Methodology, Investigation, Formal Analysis, Writing - Original Draft. **RC:** Conceptualization, Methodology, Formal Analysis, Writing - Original Draft. **MR:** Conceptualization, Methodology, Formal Analysis, Writing - Review & Editing, Supervision, Funding Acquisition.

## Acknowledgements

We thank the following individuals for assistance with data collection: Max Bluestone, Emily Iannazzi, Ari Khoudary, Samantha Murphy, and Jenna Phelan.

